# MIM-CyCIF: Masked Imaging Modeling for Enhancing Cyclic Immunofluorescence (CyCIF) with Panel Reduction and Imputation

**DOI:** 10.1101/2023.05.10.540265

**Authors:** Zachary Sims, Gordon B. Mills, Young Hwan Chang

## Abstract

CyCIF quantifies multiple biomarkers, but panel capacity is compromised by technical challenges including tissue loss. We propose a computational panel reduction, inferring surrogate CyCIF data from a subset of biomarkers. Our model reconstructs the information content from 25 markers using only 9 markers, learning co-expression and morphological patterns. We demonstrate strong correlations in predictions and generalizability across breast and colorectal cancer tissue microarrays, illustrating broader applicability to diverse tissue types.

## MAIN

Emerging Multiplexed Tissue Imaging (MTI) platforms^1–5^ produce rich, spatially resolved protein expression information that enables analysis of tissue samples at subcellular resolution^6,7^. However, the broad application of existing MTI platforms in cancer research and clinical diagnosis is hindered by high material costs, data storage requirements, and the need for specialized equipment and technical expertise to mitigate experimental variabilities. Moreover, the number of markers within MTI panels is limited by cost and time constraints encompassing image acquisition, marker selection, and validation. Tissue degradation through repeated staining and removal cycles adds to this challenge^4^. Thus, the selection of markers for the panel becomes crucial, with the aim of interrogating a wide spectrum of cell states and phenotypes^5,8,9^.

Previous studies computationally optimized MTI panel reduction and prediction. Ternes *et al*.^*10*^ pioneered CyCIF panel reduction and imputation using a two-step approach: exploring multiple strategies for marker selection and training a multi-encoder variational autoencoder (ME-VAE)^6^ to reconstruct the full 25-plex CyCIF images at the single cell level. Wu *et al*. proposed a three-step method using a concrete autoencoder and convolution neural network to reduce CODEX markers and predict intensity via a linear regression model^11^. Sun *et al*. iteratively trained a U-Net to reconstruct patch-level images, aiding marker selection for a reduced panel^12^. In contrast to prior research, where panel selection is separate from full panel reconstruction, our method integrates iterative marker selection within a pre-trained model, streamlining panel reduction and reconstruction. Unlike fixed-size reduced panels, our method optimizes marker selection during inference, enhancing efficiency, reliability, and practicality for model training.

Inspired by the success of masked language modeling in natural language processing^13^, the concept of masked image modeling (MIM) has gained traction in computer vision^14,15^. MIM-based models resemble denoising autoencoders^16^, utilized for data restoration and model pre-training. Nevertheless, the utilization of masked image modeling for missing data imputation tasks has been minimally investigated. Employing a self-supervised trained masked autoencoder (MAE), we reconstruct masked CyCIF image channels at the single-cell level and identify optimal reduced panel sets for complete panel reconstruction. Through the architecture and masked token prediction task outlined in **Figure 1A**, we demonstrate successful imputation of CyCIF image channels at the single-cell level through ‘channel in-painting’. Our model takes in a collection of single-cell images, each containing 25 channels representing individual CyCIF marker stains. During training, we set a fixed ratio of channels that will undergo random masking for each sample (**Figure 1B** left). Our model is then tasked with reconstructing these masked channels (**Figure 1B** right).

**Fig. 1.**
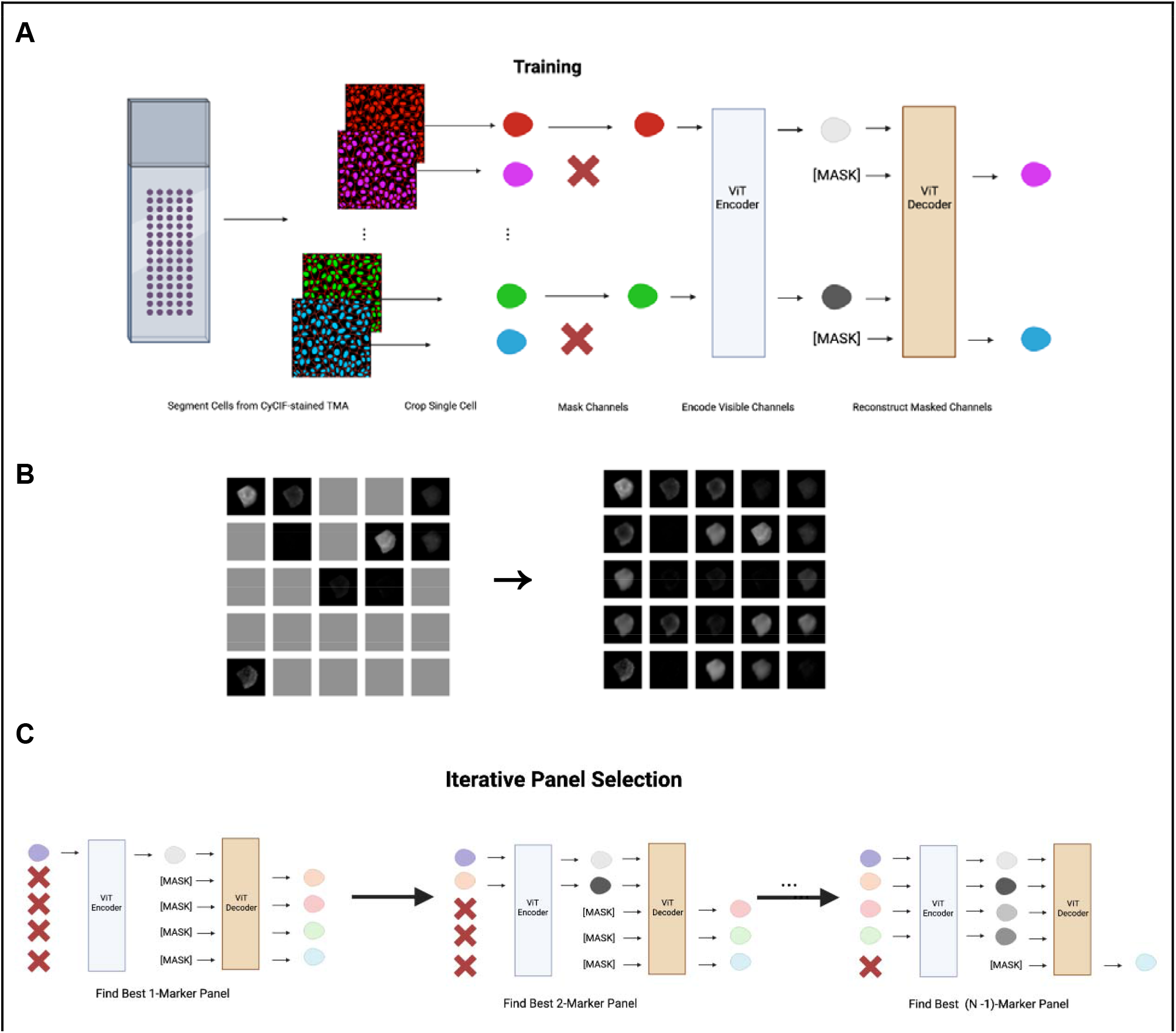
Masked autoencoder for panel reduction and marker imputation. **A. Model architecture:** CyCIF image-derived single cells undergo channel-wise masking followed by the encoding of unmasked channels using a Vision Transformer (ViT). A distinct mask token represents masked channels. A ViT decoder then reconstructs the masked channels, completing the image reconstruction process. **B. CyCIF channel-wise masking (left) and reconstruction (right)**: 25-channel images arranged into a 5 x 5 grid format, facilitating conversion from a patch-wise masking strategy into a channel-wise masking strategy. **C. Iterative marker selection:** leveraging the trained model, an optimal marker order is established by gradually increasing the panel size. Each step selects the next marker based on its ability to maximize the Spearman correlation between actual and predicted mean intensity for masked channels. This refines marker panel ordering, enhancing prediction accuracy.

After model training, the selection of masked channels becomes feasible to determine the optimal unmasked channel combination for accurate reconstruction of masked counterparts. This strategy is harnessed to progressively curate an enhanced marker panel (**Figure 1C**). In each iteration, the marker selection is the one maximizing the Spearman correlation between actual and predicted mean intensities for the remaining held-out markers (**Methods: Iterative Panel Selection**). Subsequently, this iterative process of refining panels establishes an order of markers that roughly represents predictive value regarding other markers (**Figure 2A**). Visualized in columns, the chosen markers’ influence on predicting withheld markers is depicted, while rows illustrate the corresponding improvements in prediction for each withheld marker. We additionally assess the structural similarity index measure (SSIM) (**Methods: Model’s Performance Evaluation**), a widely adopted metric capturing image similarity as perceived by the human visual system^17^, between real and predicted single-cell expression at the pixel level (**Supplementary Figure 1**).

**Fig. 2.**
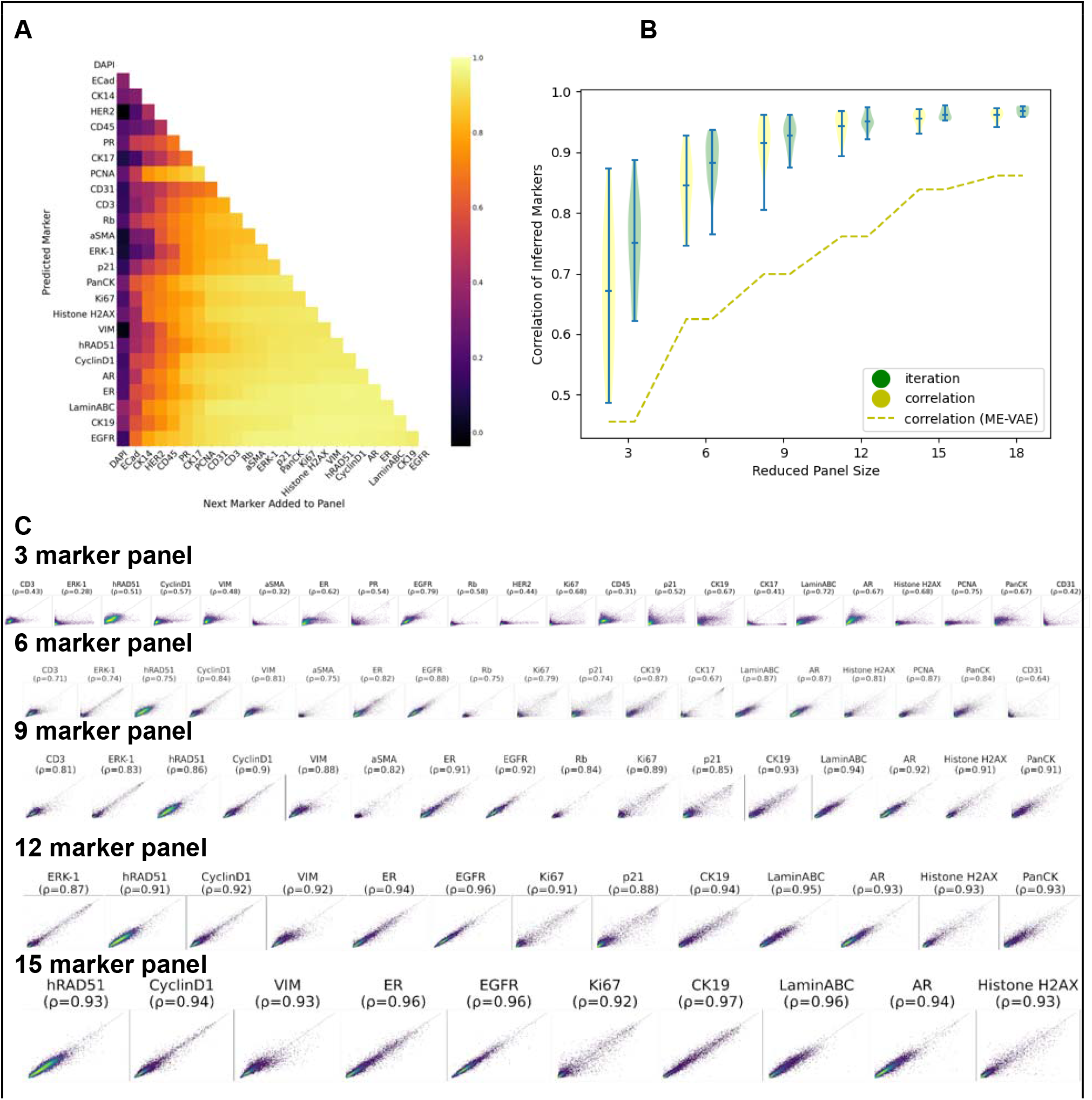
Model Evaluation. **A. Impact of individual markers:** Depicting the effect of marker selection on the prediction of specific marker intensities. Each row tracks the improvement in prediction for a specific marker as new markers are added to the reduced panel. **B. Comparison to prior work**^10^. The yellow dashed line shows the mean Spearman correlation achieved for predicted marker intensities utilizing ME-VAE. The corresponding yellow violin plot demonstrates the performance of MAE on the same reduced panels that showed the optimal results in Ternes *et al*.^*10*^ Green violin plot showcases MAE performance using reduced panels selected using the iterative panel selection approach. **C. Real versus predicted single-cell mean intensity values**. Plots of actual versus predicted single-cell mean intensity values are presented for reduced panel sizes of 3,6,9,12, and 15 markers, respectively. A random subset of 10,000 cells is shown. The Spearman correlation for each marker is indicated.

In our prior study^10^, we pre-selected optimally reduced panels comprising 3, 6, 9, 12, 15, and 18 markers through Spearman Correlation. The subset of markers exhibiting the highest correlation with the remaining withheld markers was chosen as the reduced panel. **Figure 2B** illustrates the enhancement in prediction resulting from the replacement of ME-VAE with MAE. With the same 9-marker reduced panel, MAE improves the average correlation of withheld marker predictions by 0.22 (yellow violin plot). A further enhancement is achieved by utilizing the reduced panel generated via iterative selection, yielding an additional 0.01 improvement (green violin plot). We also demonstrate similar effectiveness on the colorectal cancer (CRC) TMA dataset (**Supplementary Figures 2 and 3**).

Moreover, the model generalizes well to unseen data, demonstrated by conducting 5-fold cross-validation across the CRC TMA cores (**Supplementary Figure 4**). Additionally, we tested our model trained on the CRC TMA on a full Whole-Slide Image (WSI) stained with the same panel but in different batches (**Supplementary Figure 5**). Although the model exhibits a modest performance reduction (0.26 reduction in Spearman correlation using the same 9 marker panel), these results hold promise, particularly given potential limitations in generalizability stemming from ROI selection bias and a single TMA reference for batch correction. This finding aligns with prior findings on small TMA cores, emphasizing the enhanced representation provided by randomly selected multiple TMA cores in capturing tumor or immune contexture as compared to WSI^18,19^.

Our study highlights the efficacy of utilizing MAE to generate high-plex CyCIF data from only a few experimental measurements, significantly reducing the required biomarkers to interrogate a sample. The ability to identify a biomarker subset and perform *in silico* prediction offers several advantages. Our method empowers users to access a more extensive set of biomarkers beyond those experimentally measured. Additionally, it enables the allocation of resources for the exploration of novel biomarkers, thereby enhancing cell type differentiation and disease characterization. Furthermore, it can manage instances of assay failures such as low-quality markers, technical noise, and/or potential tissue loss in later CyCIF rounds. It also has the ability to artificially up-sample and incorporate additional panel markers. In future work, we will explore different normalization strategies^20–22^ to reduce marker intensity variability across batches. Additionally, we will explore a more diverse training dataset, incorporating WSIs in different batches, effectively mitigating TMA sampling bias and batch effects.

## METHODS

### CyCIF Image Dataset

Our data consists of CyCIF images from two TMA datasets, one containing breast cancer (BC) tissue and the other containing colorectal cancer (CRC) tissue. The tissue microarrays (TMAs) CyCIF imaging data are available via HTAN (https://humantumoratlas.org/). The breast cancer (BC) TMA contains 88 cores representing 6 cancer subtypes. The colorectal cancer (CRC) TMA contains 332 cores. Biomarker panels for the two TMAs are shown in **Supplementary Table 1**.

### Image Preprocessing

The original CyCIF marker intensities (16 bit) were rescaled to the 8 bit image ([0,255] range). For the BC TMA, we simply used preprocessed data in our previous work^10^. For the CRC TMA, any core containing a channel with a mean intensity beyond 2 standard deviations from the mean intensity of the entire TMA for that channel was dropped. This resulted in 12 cores being removed. Individual cores were then segmented using MESMER^23^. We use the whole cell masks generated using the max projection of the PanCK and CD45 channels as the membrane marker to crop each cell down to a 32x32 pixel region. The background of each single-cell image is then zeroed out, and the polar axis and center of mass are aligned. This resulted in 742,169 cells for the CRC TMA and 691,893 cells for the BC TMA. We use 90% of the cells randomly selected as the training set, and withhold 5% for validation/panel selection and 5% for testing.

### Masked Image Modeling

We modify the patch-wise masking strategy in MAE^14^ to a channel-wise masking strategy. This involves resizing the 32x32x25 multichannel single-cell image (32x32) to a 5 x 5 grid format (**Figure 1B**), creating a resulting image size of 160x160 (i.e., (32x5)x(32x5)) with a single channel. Consequently, the patch size for MAE is adjusted to 32x32, where each patch now corresponds to an individual channel within the original multi-channel image.

We use a Vision Transformer (ViT) encoder and a ViT decoder as in MAE, both set to 8 heads and 6 layers, and a 2048-dimension multilayer perceptron layer. The embedding dimensions were 1024 for the encoder and 512 for the decoder. We train the model using 8 Nvidia A40 GPUs for 300 epochs using a batch size of 512, Adam optimizer and a learning rate of 1e-3.

Although the trained model works on a range of reduced panel sizes, during training the number of channels to be masked is set to a fixed ratio. We evaluate different masking ratios for training by assessing the performance of different reduced panel sizes in inference on the BC TMA dataset. For testing, we choose the optimal reduced panels identified in Ternes *et al*.^*10*^, which have sizes of 3,6,9,12,15, and 18 markers (88%, 76%, 64%, 52%, 40%, and 28% masking ratios, respectively). We train three models using a fixed masking ratio of 25%, 50%, and 75%, and find that the 50% masking ratio results in the best overall performance across different panel sizes in inference (**Supplementary Figure 6**).

### Iterative Panel Selection

To obtain optimal reduced panel sets, we leverage the trained model to determine which markers are most informative. To do this, we iteratively determine an ordering of markers such that the first □ markers result in the best reconstruction of the remaining □ − □ markers, measured by the Spearman correlation of the predicted mean intensity at the single cell level. We start with □ = 2, setting the first marker in the order to be DAPI, as nuclear staining is important for downstream analysis such as registration as well as determining cell morphology. We then iterate through the remaining 24 markers to determine which marker, along with DAPI, results in the best reconstruction of the remaining 24 markers (**Figure 1C** and **2A**). We repeat this process until we find the best marker panel:

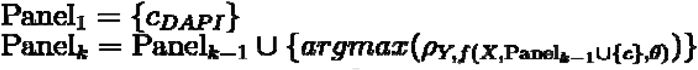

Where is the marker channel being considered for inclusion into **Panel**_***k***_,***Y***, is the set of ground truth masked channels, ***X*** is the set of unmasked channels, ***f*** is the trained MAE model parameterized by ***θ***, which returns the reconstructed masked channels, and ***ρ***is the Spearman correlation between the mean intensities of ***Y*** and the output of ***f***.

### Model’s Performance Evaluation

We evaluate the model’s performance using Spearman correlation and SSIM. We assess the agreement between actual and the predicted mean marker intensities within the cell boundaries. This analysis is crucial as it quantifies specific cellular component expression levels across markers, characterizing cellular phenotypes. We avoid directly comparing and classifying cell types in subsequent analysis to avoid oversimplifying complex cellular phenotypes. It is important to note that cell type determination in MTI settings involves diverse methodologies and can be influenced by factors like imperfect cell segmentation, marker selection, data preprocessing and normalization^20–22^, and algorithm choice^24^.

### 5-Fold Cross-validation

Cross-validation was performed on the CRC TMA dataset to evaluate model performance on unseen data. We divide the dataset at the TMA core level by separating the cores into 5 sets of 64 TMA cores for the test sets and the remaining TMA cores are used to train 5 separate models. As TMAs typically encompass multiple patients, this test effectively demonstrates the model’s generalizability. The performance of the 5 models on different reduced panel sizes is shown in **Supplementary Figure 4**.

### WSI Testing and Batch Effect

To further demonstrate model generalizability, we test the CRC model on a whole-slide image (WSI) stained using the same panel. Segmentation produced a dataset consisting of 742,799 cells in the WSI. To mitigate batch-to-batch staining variations, the intensity distributions for each channel in the WSI were normalized using histogram matching with 3 cores from the CRC TMA-obtained from the same tissue section as the WSI. Although this approach allows us to address batch effects while preserving intra-patient intensity distributions, a potential limitation is sample bias introduced during TMA core selection. A histopathologist would heavily favor tumor regions for core punchouts, whereas the full WSI contains a more heterogeneous tissue region. Therefore, markers that are not expressed in tumor cells are potentially underrepresented in our training set. **Supplementary Figure 5** shows the performance of the model, trained on the CRC TMA with reduced panels selected from the TMA, on this dataset.

## Data Availability

As part of this paper all images at full resolution, all derived image data (e.g. segmentation masks), and all cell count tables will be publicly released via the NCI-recognized repository for Human Tumor Atlas Network (HTAN; https://humantumoratlas.org/) at Sage Synapse. Note to reviewers: this data resource is undergoing final review for Private Health Information and requires a (free) Synapse account; public access will be provided as soon as possible. An anonymous “reviewer only” link can be provided prior to that by requesting it from the monitoring editor.

## Code Availability

All software used in this manuscript is detailed in the article’s Methods section and its Supplementary Information. The associated scripts are freely available via GitHub as described at https://github.com/zacsims/IF_panel_reduction.

## Acknowledgments

We thank Jerry Lin, Yu-An Chen, and Peter K. Sorger (Harvard Medical School) for sharing data and providing useful feedback. This work was carried out with major support from National Cancer Institute (NCI) Human Tumor Atlas Network (HTAN) Research Centers at OHSU (U2CCA233280). Y.H.C is supported by R01 CA253860 and Kuni Foundation Imagination Grants. The resources of the Exacloud high-performance computing environment developed jointly by OHSU and Intel and the technical support of the OHSU Advanced Computing Center are gratefully acknowledged.

## Supplementary Information

**Supplementary Fig. 1.**
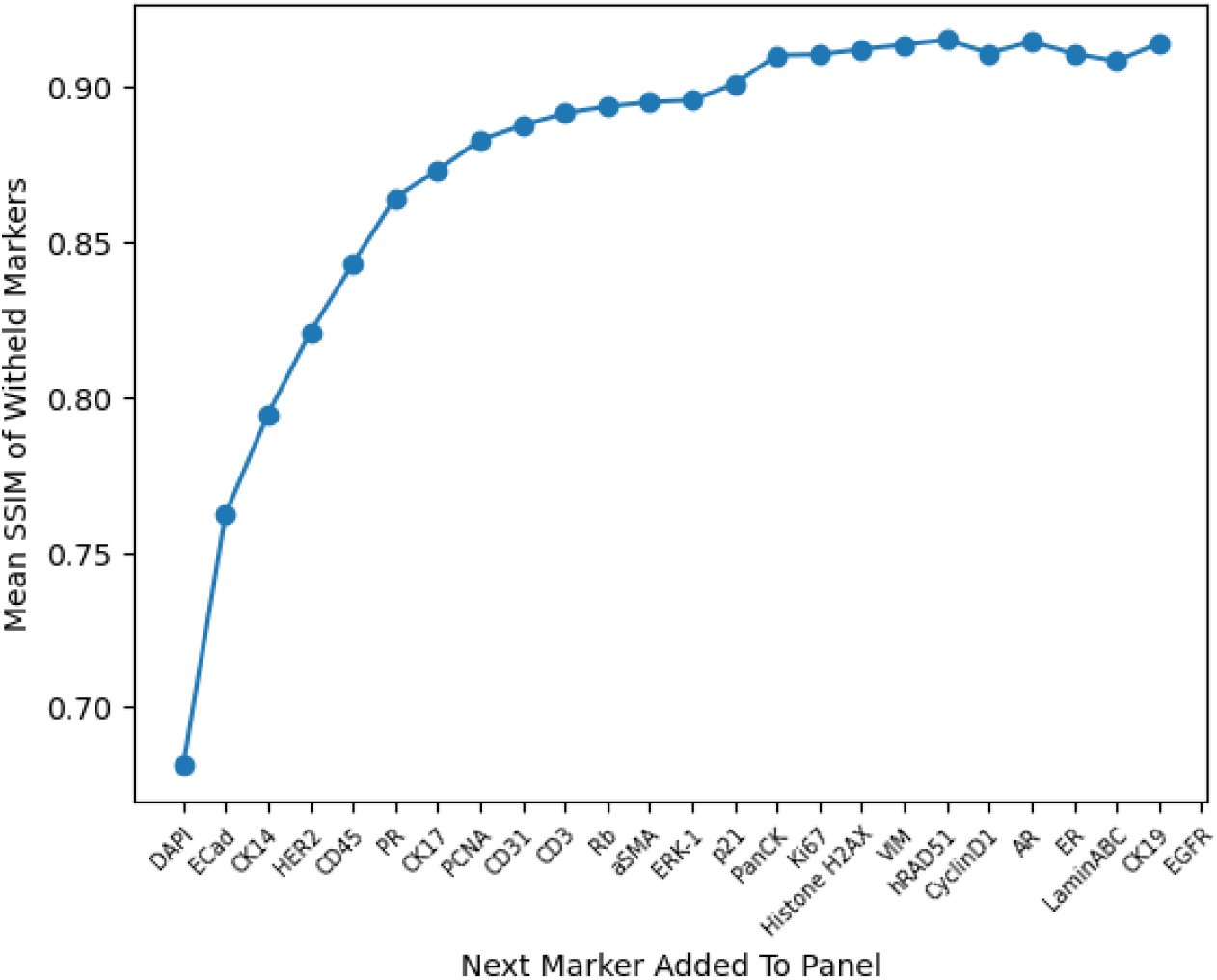
Structural Similarity Index. The structural similarity index for the BC TMA.

**Supplementary Fig. 2.**
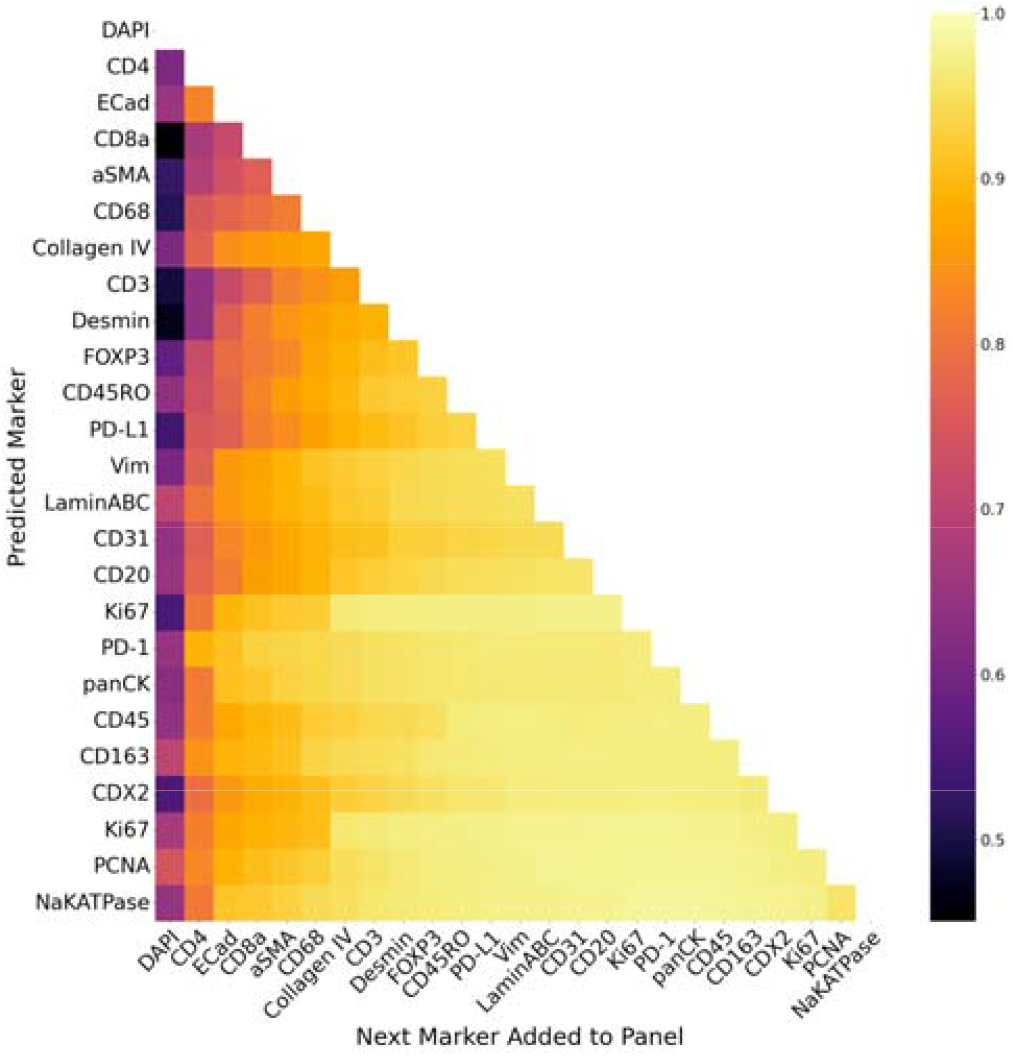
CRC Panel selection and marker prediction. The ordering of markers in the CRC panel, determined by iterative selection, and the correlation for each predicted marker.

**Supplementary Fig. 3.**
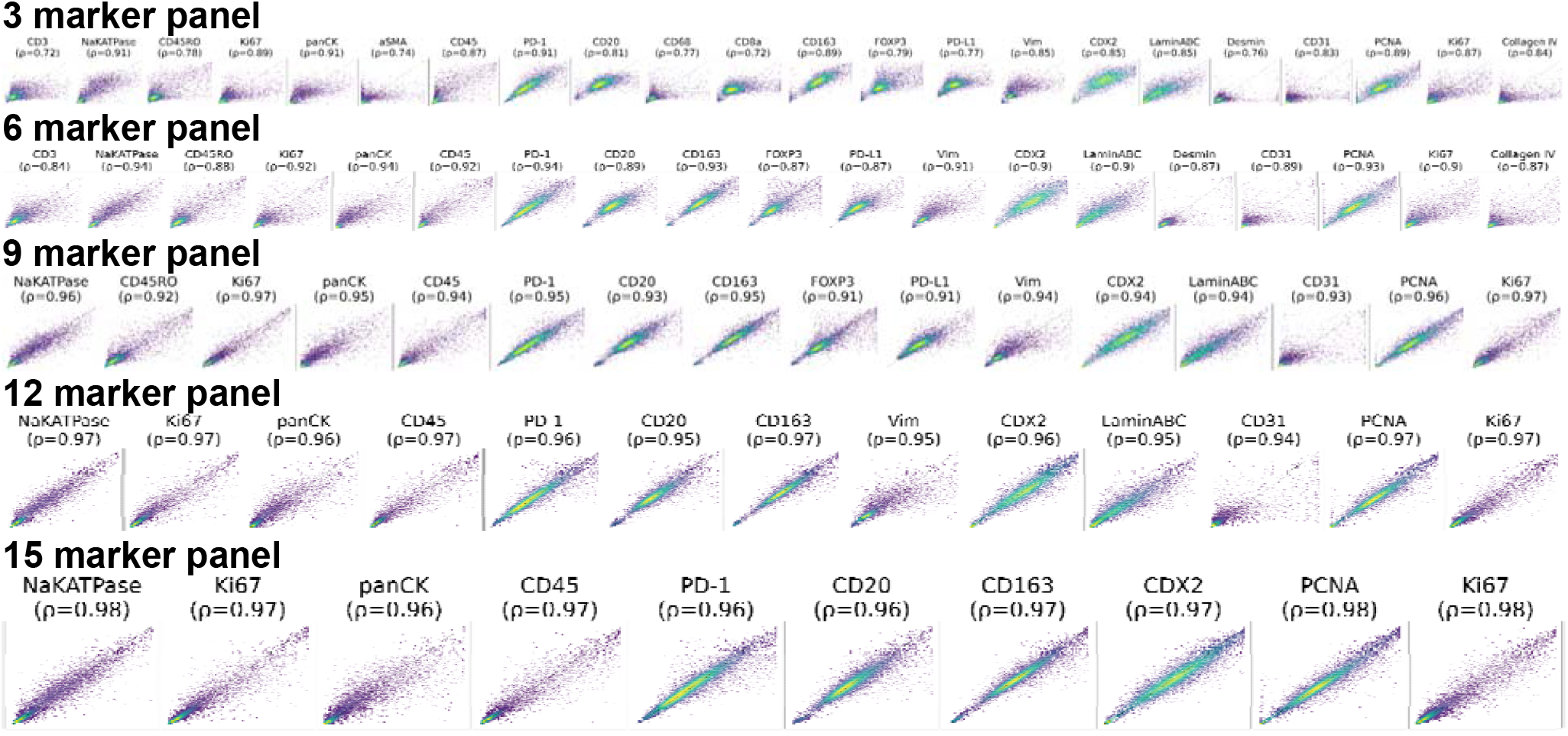
CRC marker prediction. Real versus predicted single-cell mean intensity values. Plots of actual versus predicted single-cell mean intensity values are presented for reduced panel size of 3,6,9,12, and 15 markers, respectively. A random subset of 10,000 cells is shown. The Spearman correlation for each marker is indicated.

**Supplementary Fig. 4.**
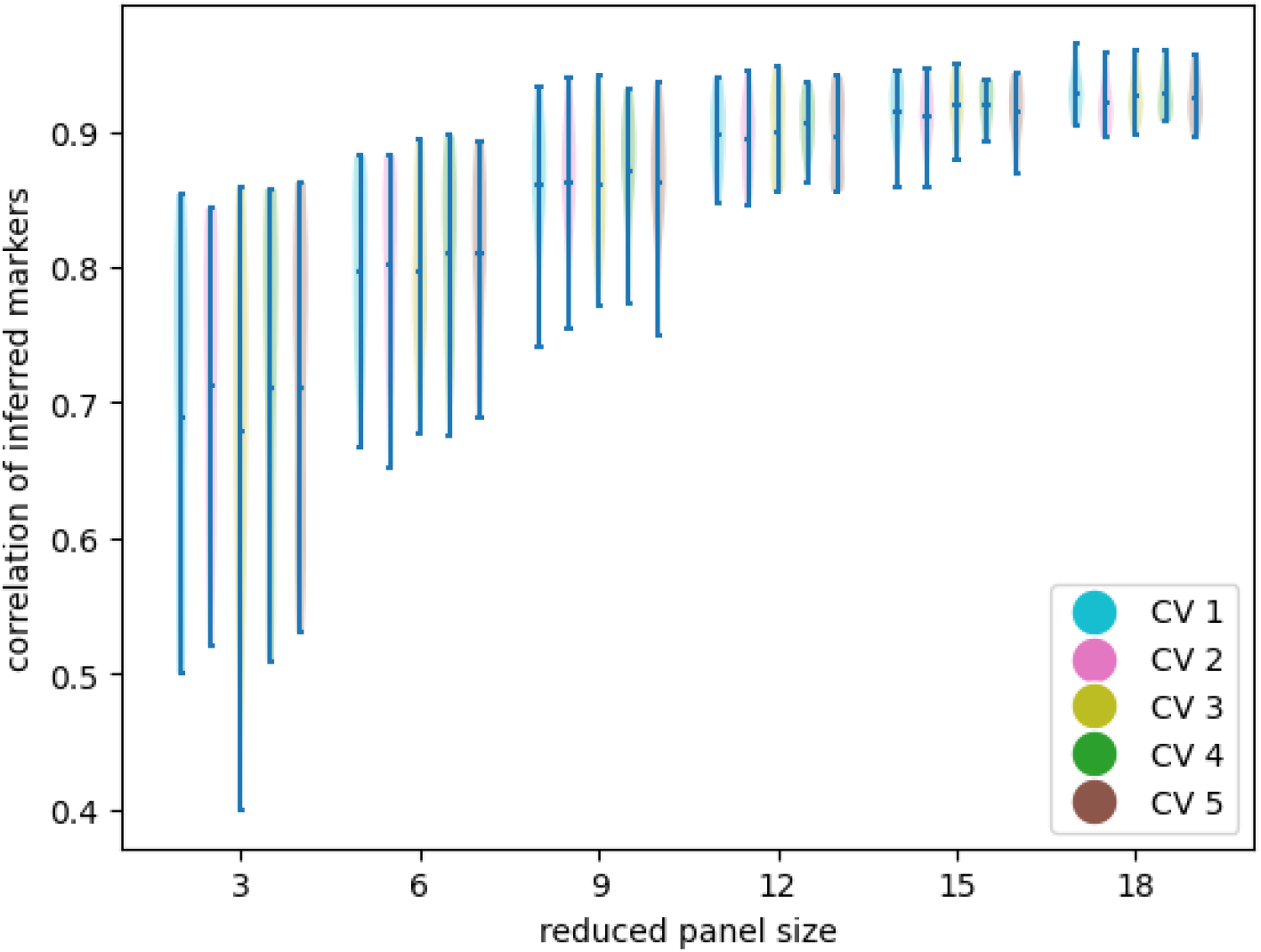
Cross-validation results for CRC TMA analysis across six distinct reduced panel sizes. Cross-validation results, employing a 5-fold approach on the CRC TMA for six different panel sizes (3,6,9,12,15 and 18 markers), highlights the robustness and generalizability of the panel reduction over various withheld datasets.

**Supplementary Fig. 5.**
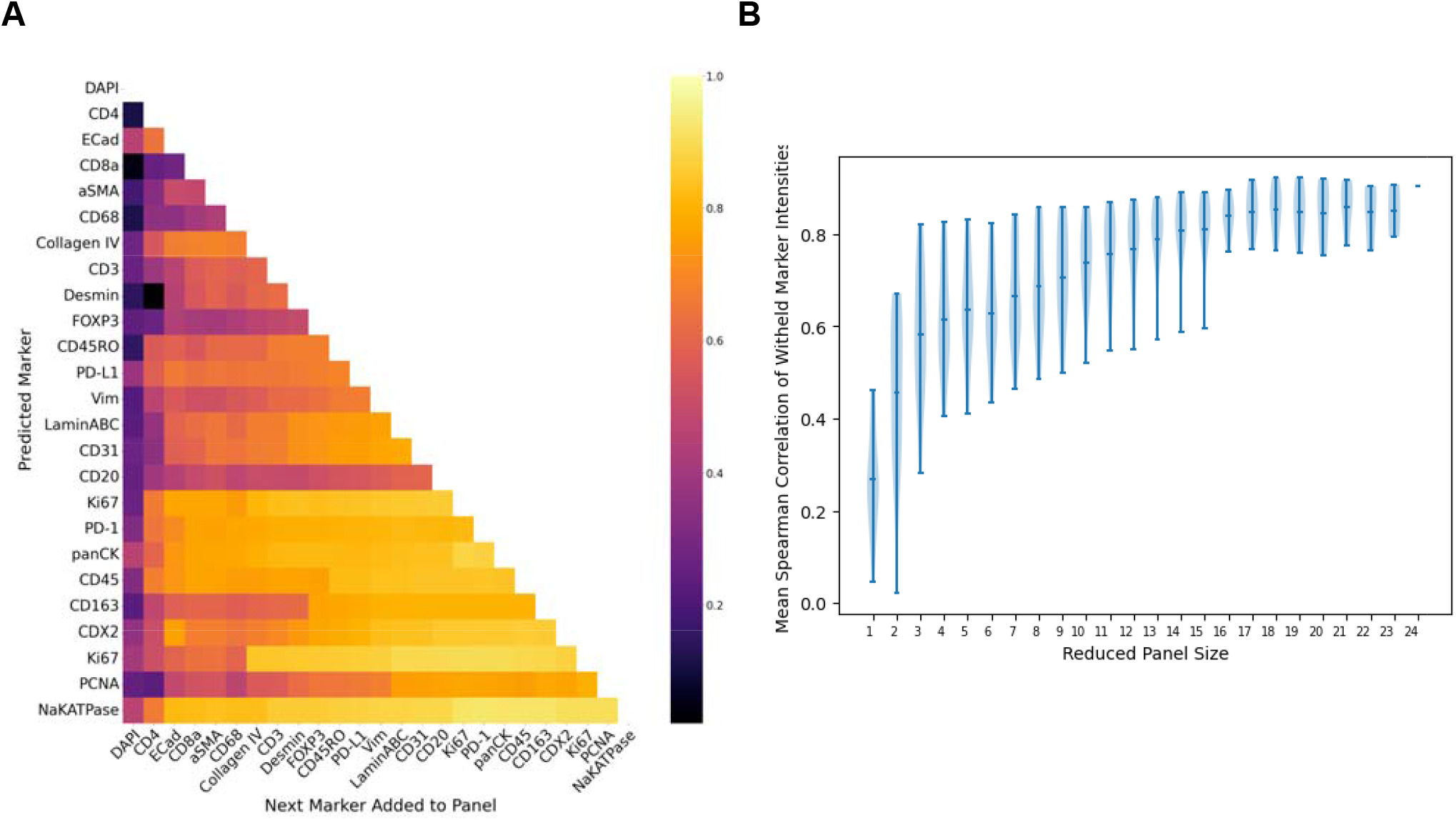
Prediction Result for WSI Test Set. **A.** Correlations for predicted markers on the CRC WSI dataset using reduced panels selected from the CRC TMA dataset. **B**. Performance of the pre-trained model on different reduced panel sizes.

**Supplementary Fig. 6.**
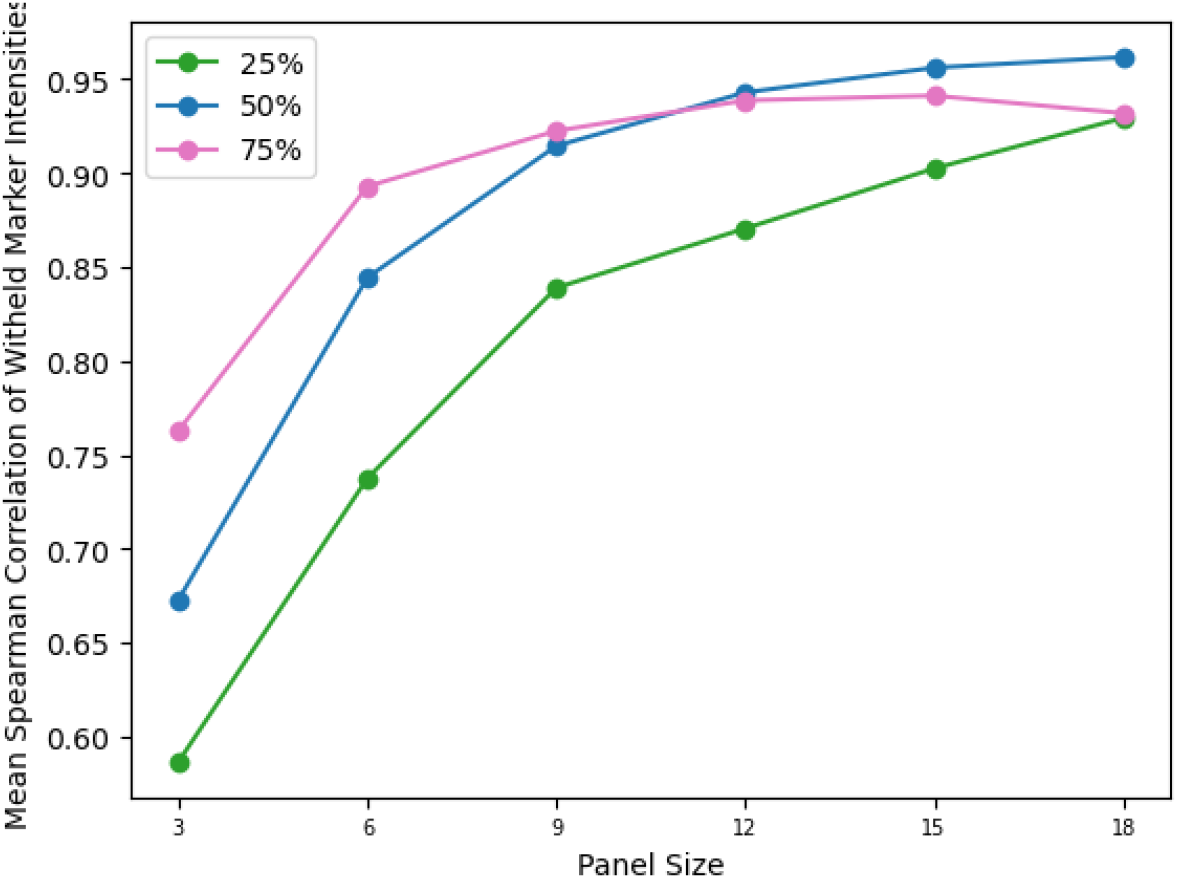
Masking Ratio Evaluation in BC TMA. The ratio of masked channels is fixed during training. Models trained with different ratios are compared by measuring the performance on different masking ratios in inference. The model trained at a 50% masking ratio performs the best across different masking ratios in inference.

**Supplementary Table. 1.** CyCIF Biomarker Panels.

|  | Breast Cancer CyCIF Panel |  |  |  | Colorectal Cancer CyCIF Panel |  |  |  |
| --- | --- | --- | --- | --- | --- | --- | --- | --- |
| Cycle 1 | DAPI | CD3 | ERK-1 | hRAD51 | DAPI | CD3 | NaKATPase | CD45RO |
| Cycle 2 | DAPI | CyclinD1 | Vim | aSMA | DAPI | Ki67 | PanCK | aSMA |
| Cycle 3 | DAPI | ECad | ER | PR | DAPI | CD4 | CD45 | PD-1 |
| Cycle 4 | DAPI | EGFR | Rb | HER2 | DAPI | CD20 | CD68 | CD8a |
| Cycle 5 | DAPI | Ki67 | CD45 | p21 | DAPI | CD163 | FOXP3 | PD-L1 |
| Cycle 6 | DAPI | CK14 | CK19 | CK17 | DAPI | ECad | Vim | CDX2 |
| Cycle 7 | DAPI | LaminABC | AR | Histone H2AX | DAPI | LaminABC | Desmin | CD31 |
| Cycle 8 | DAPI | PCNA | PanCK | CD31 | DAPI | PCNA | Ki67 | Collagen IV |

## Notes

### Competing Interest Statement

The authors have declared no competing interest.

### Summary of Updates

We demonstrate strong correlations in predictions and generalizability across breast and colorectal cancer tissue microarrays, illustrating broader applicability to diverse tissue types.

